# Gαq-stimulated gene expression is insensitive to Bromo extra terminal domain inhibitors in HEK 293 cells

**DOI:** 10.1101/2025.06.23.661130

**Authors:** Ashika Jain, Viviane Pagé, Dominic Devost, Darlaine Pétrin, Terence E. Hébert, Jason C. Tanny

## Abstract

*Bromodomain and extraterminal domain* (BET) family proteins are ubiquitous transcriptional co-activators that function broadly in gene expression programs associated with cellular differentiation, proliferation, and stress responses. Pharmacological inhibition of BET proteins with small molecules that disrupt bromodomain engagement with acetyllysine residues (such as JQ1), or that drive their degradation through the ubiquitin-proteasome system (such as dBET6), ameliorates pathological gene expression in a range of systems and shows promise as a potential therapeutic strategy. BET involvement in gene expression responses is not universal, and understanding of the cell-type and signaling pathway requirements that dictate BET dependence remains incomplete. We previously demonstrated that, in neonatal rat cardiomyocytes, GPCR-induced hypertrophy response depended strongly on the BET protein Brd4 when signaling was coupled to Gαs, but not Gαq. Here we tested whether Brd4 was differentially responsive to G protein isoforms in HEK 293 cells by expressing Gαs or Gαq-coupled *D*esigner *R*eceptors *E*xclusively *A*ctivated by *D*esigner *D*rugs (DREADDs). Gαq induced expression of a group of early response genes and inflammatory genes in a manner largely insensitive to pharmacological BET inhibition, consistent with our previous data in cardiomyocytes. Gαs activated a small subset of the Gαq-induced genes, but this effect was largely reversed by the BET degrader molecule dBET6. Our data further suggest that there may be general signaling requirements to activate Brd4 across cell types.

**Highlights:**

- We tested whether Bromodomain and extraterminal domain (BET) family proteins were differentially responsive to G protein isoforms in HEK 293 cells by expressing Gαs or Gαq-coupled *D*esigner *R*eceptors *E*xclusively *A*ctivated by *D*esigner *D*rugs (DREADDs).
- Gαq induced expression of a group of genes but was largely insensitive to pharmacological BET inhibition, consistent with our previous data in cardiomyocytes.
- Gαs activated a small subset of the Gαq-induced genes, but was sensitive to the BET degrader molecule dBET6.
- Distinct G protein-dependent effects may be conserved across cell types.

## 1. Introduction

The bromodomain and extraterminal domain (BET) family of transcriptional co-activators are integral components of both physiological and pathological gene expression programs. BET proteins have been studied extensively as drivers of cancer cell proliferation, both for hematological malignancies and solid tumors [1,2]. BET proteins are also required for inflammatory gene expression, pathological cardiac hypertrophy, and spermatogenesis [3–8]. The BET family consists of Brd2, Brd3, and Brd4, all of which are broadly expressed in mammalian tissues, as well as the testis-specific variant BrdT. Brd4 is thought to be the relevant player in most of the functions mentioned above [1,2]. All BET proteins contain two tandem bromodomains, which interact with acetyllysines on histones and other regulatory proteins [1]. The extraterminal domain binds to multiple chromatin-modifying enzymes, although the regulatory significance of these interactions is not understood [9,10]. A C-terminal domain in Brd4 and Brdt binds to the positive transcription elongation factor b complex P-TEFb [11]. Brd4 also associates with the Mediator complex and participates in recruitment of RNA polymerase II to promoters and enhancers throughout the genome [10,12,13]. Thus, BET proteins, and Brd4 in particular, act broadly to affect multiple steps in RNAPII transcription.

Interest in BET proteins has been spurred by the development of small molecules that bind to the BET bromodomains, thereby competing with acetyllysine and diminishing BET chromatin occupancy. These BET inhibitors (such as JQ1, iBET-151, or RVX-208) have shown beneficial effects in cell culture and animal models of multiple types of cancer, inflammatory disease, and cardiac hypertrophy [14–18]. More recently, these have been converted into effective BET degraders (such as dBET6) through chemical linkage to small molecules that bind to the VHL or cereblon E3 ubiquitin ligases [19,20]. It is critical to further define how BET proteins act in transcription to guide future use of these potential therapeutics.

Despite their connections to multiple generally acting components of the RNAPII transcriptional machinery, BET proteins have gene-selective roles in transcription that are cell-type specific and remain poorly understood. Studies in cancer cell lines have demonstrated that sensitivity of particular cell lines to the growth restricting effects of BET inhibitors correlated with the activity of super enhancers proximal to key oncogenic driver genes (such as c-myc)[1,21,22]. As originally defined, super enhancers are distinguished from conventional enhancers by virtue of their large size (often 20 kb in length or more), as well as their association with exceptionally high levels of Mediator and Brd4 [22–24]. However, it is unclear whether similarly designated super enhancers are important for other transcriptional responses that require Brd4 [25–27]. Brd4 action at specific promoters or enhancers may instead be dictated by interactions with DNA-binding transcription factors [28–30]. BETs are also regulated by post-translational modification. Brd4 is hyperphosphorylated in several cancer cell lines, and phosphorylation by multiple cyclin-dependent kinases or by casein kinase II has been associated with enhanced chromatin binding [28,31,32]. Such a mechanism is also implicated in Brd4-dependent activation of immediate early genes in cortical neurons [33]. Cytokine signaling leads to JAK2-dependent phosphorylation of Brd4 in colorectal cancer cells, thereby preventing Brd4 protein degradation by the ubiquitin proteasome system and enhancing transcription of key oncogenic genes [34]. BET protein stability is regulated by polyubiquitylation in multiple cell types; altered regulation of these pathways in cancers have been associated with resistance to BET inhibitors [35–37]. Thus, BET proteins are dynamic regulators that are highly responsive to inputs from cellular signaling pathways.

We previously demonstrated that in a cellular model of neonatal rat cardiomyocyte hypertrophy, Brd4 activity was differentially sensitive to two hypertrophic stimuli acting through different G protein-coupled receptors (GPCRs)[38]. If hypertrophy was induced by activation of the endothelin A (ET_A_) receptor using the ligand endothelin 1, it proceeded relatively unimpeded by the BET inhibitor JQ1. However, stimulation of hypertrophy through activation of the α1-adrenergic receptor with the agonist phenylephrine was JQ1-sensitive. Concordantly, Brd4 occupancy at promoters and enhancers of hypertrophy-induced genes was enhanced by phenylephrine, but not by endothelin-1. These differences appeared to be due to the distinct signaling pathways activated by the two receptors: The ET_A_ receptor is coupled to the Gαq G protein isoform, whereas the α1-adrenergic receptor is coupled to Gαq as well as Gαs [38,39]. Gαs-dependent signaling appeared to be important for Brd4 activation, as Brd4 occupancy in chromatin immunoprecipitation (ChIP) experiments could be enhanced by stimulating adenylate cyclase or blocked by inhibition of protein kinase A (PKA) [38].

Based on these results, we hypothesized that Brd4 participation in GPCR-driven transcriptional responses would necessitate signaling through Gαs, and that responses following Gαq signaling would be relatively Brd4-independent. We sought to test this using a tractable model cell line and engineered GPCRs that couple to defined G protein subtypes. The results were consistent with our hypothesis and may help to define cellular contexts that dictate responsiveness to BET inhibitors.

## 2. Materials and Methods

### 2.1 Cell culture and drugs

HEK 293(PL) cells were grown in Dulbecco’s Modified Eagle’s medium (DMEM) high glucose + 5% (v/v) fetal bovine serum + 1% (v/v) penicillin/streptomycin at 37°C. JQ1 (Abcam, ab141498) and dBET6 (Sigma-Aldrich, SML2683) were dissolved in dimethyl sulfoxide (DMSO) at 5 mM and 500 μM, respectively. Deschloroclozapine (DCZ; Hellobio, HB9126) was dissolved in sterile ddH_2_O at 10 mM.

### 2.2 Plasmids

Plasmids for expression of the Gαq-DREADD (pcDNA5/FRT-HA-hM3D) and the Gαs-DREADD (pcDNA5/FRT-HA-rM3D) were obtained from Addgene [40,41]. Plasmids for expression of the PKC and EPAC biosensors have been described previously [39]. Transfection into HEK 293 cells was performed with Lipofectamine 2000 (Invitrogen) as per the manufacturer’s instructions.

### 2.3 Bioluminescence resonance transfer (BRET) biosensor assays

Cells were transfected with the Gαq-DREADD, the Gαs-DREADD, or a vector control (pcDNA 3.1; [39]) in combination with either a PKC biosensor plasmid or a EPAC biosensor plasmid. After transfection, cells were plated at a density of 30,000 cells/well in a poly-L-ornithine-coated 96-well white bottom plate (Thermo Scientific, 236105) and incubated for 24 hours. Media was removed from each well, cells were washed with 150 μl Krebs buffer, and 80 μl Krebs buffer was added to each well. Fluorescent substrate coelenterazine 400A (Cedarlane; 10 μl) was added to each well. BRET measurements were performed in a TriStar 2 Multimode Plate Reader (Berthold Technologies) as described previously [42]. Briefly, the plate was maintained at 28°C for one hour, the last 5 minutes of which were used to take a basal reading in the 410-515 nm spectral range. DCZ or vehicle was then added, and further readings taken at 28°C after the indicated time intervals.

### 2.4 BRET analysis

BRET ratios were determined by calculating the ratio of the light emitted by GFP10 over the light emitted by the RlucII, and results were expressed as ΔBRET = BRETagonist – BRETbasal [43]. ΔBRET was calculated as follows:

*ΔBRET_Gαs_ = [(Gαs-DREADD-EPAC/DCZ)–(Gαs-DREADD-EPAC)] – [(pcDNA-EPAC/DCZ) – (pcDNA-EPAC)]*.

Δ*BRET_Gαq_ = [(G*α*q-DREADD-PKC/DCZ)–(G*α*q-DREADD-PKC)] – [(pcDNA-PKC/DCZ)– (pcDNA-PKC)]*.

An average of three technical replicates were used for each measurement. Data were plotted as ΔBRET versus log [DCZ] with GraphPad Prism 10.0.

### 2.4 Protein extraction and immunoblotting

Cells were seeded at 350,000 per well in 6-well plates. Following a 24-hour incubation, cells were treated with 100 nM dBET6 or vehicle, washed twice with cold phosphate-buffered saline (PBS), and suspended in 200-400 μl cold RIPA buffer [1% NP-40, 50 mM Tris-HCl pH 7.4, 150 mM NaCl, 1 mM EDTA, 1 mM EGTA, 0.1% SDS, 0.5% sodium deoxycholate, 1 mM PMSF, protease inhibitor tablets (Millipore Sigma)]. Suspensions were transferred to 1.5 mL tubes, mixed, and incubated for 15 minutes on ice. Lysates were centrifuged at 14,000 x g for 15 mins at 4°C. Protein concentrations of the supernatants were measured using a BCA kit (Thermo Scientific) as per the manufacturer’s instructions. Thirty μg of supernatant was subjected to SDS-PAGE on an 8% gel, and the gel was transferred to a nitrocellulose membrane as described previously [38]. Immunoblotting was performed with anti-BRD4 (Invitrogen PA5856620), anti-β-tubulin (Invitrogen, 32-2600), or anti-GAPDH (Invitrogen, AM-4300) antibodies as described previously [38]. Blots were imaged with a GE Amersham Imager 600. Images were processed with ImageJ software.

### 2.5 RNA analysis by RT-qPCR

Cells were seeded at 350,000 per well in 6-well plates. Following a 24-hour incubation, cells were treated with 1 μM JQ1 or vehicle and washed twice with cold PBS. Cells were lysed with TRI reagent® (Sigma), and lysates were transferred to 1.5 mL tubes followed by extraction with bromochloropropane. After a 15-minute incubation at room temperature, lysates were centrifuged at 13000 x g for 15 minutes at 4°C. The aqueous phase was precipitated with an equal volume of isopropanol and RNA pellets were collected by centrifugation at 13000 x g for 8 minutes at 4°C. RNA pellets were washed with 70% ethanol and dissolved in RNAse-free ddH_2_O. Concentration of the isolated RNA was measured with a Nanodrop 2000 spectrophotometer. One μg of total RNA was used as template for cDNA synthesis with random hexamers (IDT) and M-MLV reverse transcriptase as previously described [38]. For qPCR, cDNA was diluted 1:10 and amplified with primer pairs complementary to c-myc or in GAPDH (sequences available upon request) using BrightGreen 2X qPCR Mastermix (Applied Biological Materials) and a ViiA 7 Real-Time PCR System (Thermo Scientific). The qPCR amplification curves were analyzed via the 2^-ddCt^ method as described previously [38].

### 2.6 RNA-sequencing

Cells were transfected with DREADD plasmids and seeded at 350,000 cells per well in 6-well plates. Following a 24-hour incubation, cells were treated with 100 nM dBET6, 1 μM JQ1, or vehicle for 3 hours. During the last hour of inhibitor treatment, DCZ (1 μM) or vehicle was added. Cells were washed twice with cold PBS, and total RNA was extracted using the Qiashredder Homogenization kit and the RNeasy Mini kit (Qiagen) as per the manufacturer’s instructions. RNA concentrations were measured with a Nanodrop 2000 spectrophotometer, and RNA quality was assessed on an Agilent 2100 Bioanalyzer. RNA samples (100 ng) with a RIN score >7 were submitted to Genome Quebec for paired-end library preparation and sequencing. Libraries were generated from 100 ng of total RNA and mRNA enrichment was performed using the NEBNext Poly(A) Magnetic Isolation Module (New England BioLabs). cDNA synthesis was performed with NEBNext UltraExpress RNA Library Prep Kit (New England BioLabs) as per the manufacturer’s recommendations. Libraries were quantified using the KAPA Library Quanitification Kits - Complete kit (Universal) (Kapa Biosystems). Average fragment size was determined using a Fragment Analyzer 5300 (Agilent) instrument. The libraries were normalized and pooled and then denatured in 0.02N NaOH and neutralized using pre-load buffer. The pool was loaded at 140 pM on a Illumina NovaSeq X Plus 25B lane following the manufacturer’s recommendations. The run was performed for 2×100 cycles (paired-end mode) (150 base pair reads at a depth of ∼25 million reads per sample for 24 samples). A phiX library was used as a control and mixed with libraries at 1% level. Program BCL Convert 4.2.4 was then used to demultiplex samples and generate FASTQ files.

### 2.7 RNA-seq data analysis

FASTQ files were subjected to adaptor trimming and FASTP (v0.23.4) was used to filter the low quality and duplicate reads [44]. The sequences were then aligned to the *Homo sapiens* genome (GRCh38, NCBI # GCF_000001405.26) using STAR (v2.7.11b)[45]. Gene-level read count matrices were obtained using FeatureCounts (v2.0.1)[46]. Differential expression analysis was performed with DESeq2 (v1.42.1)[47]. Pearson correlations and principal component analysis plots to compare biological replicates were generated using DESeq2. Venn diagrams comparing differentially expressed genes were generated using a web-based tool (https://bioinformatics.psb.ugent.be/webtools/Venn/). Volcano plots representing differentially expressed genes were generated using ggplot2. KEGG pathway enrichment was plotted using GOplot (v1.0.2).

### 2.8 Data availability

RNA-seq datasets were deposited in the NCBI Gene Expression Omnibus (GEO) under accession GSE299127.

## 3. Results

### 3.1 Functional validation of DREADD constructs

We used a Gαq-DREADD and a Gαs-DREADD to confer selective G protein coupling [40,41,48]. We expressed each DREADD in HEK 293 cells to interrogate the effects of these signaling pathways on gene expression. To verify that the DREADDs were functional in our system, we tested doses of deschloroclozapine (DCZ), a potent DREADD ligand, over a 5-log range in cells expressing one or the other DREADD as well as bioluminescence resonance energy transfer (BRET)-based biosensors that monitor either Gαq or Gαs-coupled signaling [39,49]. BRET was detected in cells expressing the Gαq DREADD and a PKC biosensor, but not in cells expressing the biosensor with empty vector (**Figure 1A**). Similarly, BRET was detected in cells expressing Gαs DREADD and the EPAC biosensor, but not in cells expressing the biosensor with empty vector (**Figure 1B**). The response to DCZ for either DREADD was detected over control at 100 nM at 10-, 20-, or 30-minute treatment times (**Figures 1A, 1B**). Responses appeared to peak at 1 mM for the 20 and 30-minute treatments; the 1 mM responses were sustained for up to 1 hour for both DREADDs (**Figure 1C**).

**Figure 1.**
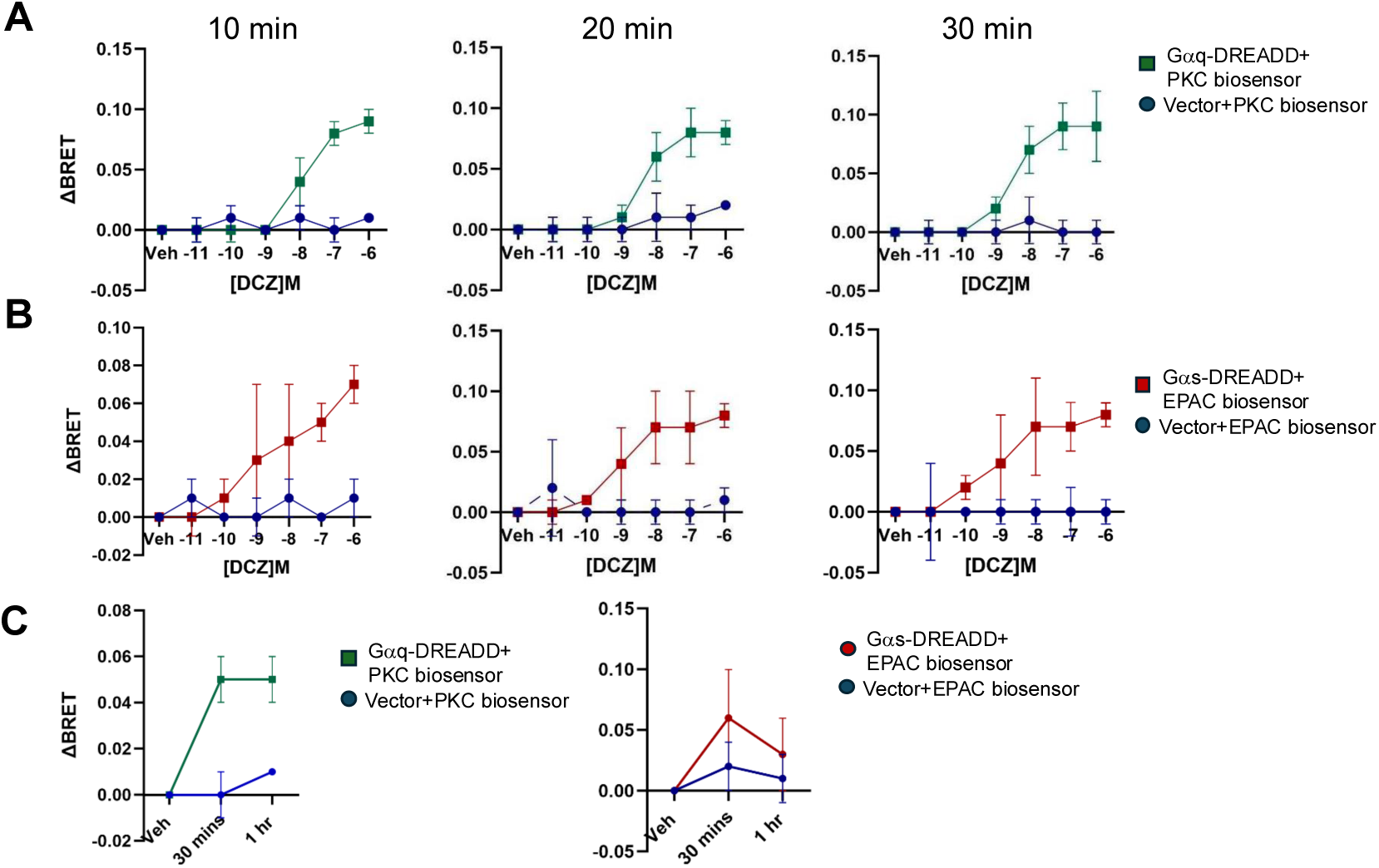
Functional validation of Gαq- and Gαs-coupled DREADDs using BRET biosensors. **(A)** ΔBRET was measured in HEK 293 cells transfected with the PKC biosensor and either Gαq-DREADD or a vector control. Measurements were taken at the indicated doses of DCZ at the indicated times (n=3 independent experiments; error bars indicate SEM). “Veh” indicates vehicle control for DCZ (ddH_2_O). **(B)** As in (A) for the EPAC biosensor and either Gαs-DREADD or vector control. **(C)** ΔBRET was measured in HEK 293 cells transfected with either DREADD/biosensor combination after 1 μM DCZ treatment for the indicated times (n=3 independent experiments; error bars indicate SEM). “Veh” indicates 1 hour vehicle control for DCZ (ddH_2_O) treatment.

### 3.2 Confirmation of BET inhibitor activity

We next verified the effectiveness of standard BET inhibitors in our system. Treatment of DREADD-transfected HEK 293 cells with 100 nM dBET6 (a dose based on previous studies; [20]) led to ∼75% depletion of Brd4 protein after 3 hours, as assessed by immunoblot (**Figure 2A**). Immunoblotting confirmed that dBET6 was similarly effective in the presence or absence of DCZ in DREADD-expressing cells (**Figure S1**). We used c-myc mRNA levels to assess the efficacy of 1mM JQ1 treatment, as c-myc transcript levels are sensitive to JQ1 in several transformed cell lines [21,50]. Unexpectedly, c-myc transcripts increased over time upon JQ1 treatment (**Figure 2B**). This pattern was not altered by the DREADDs and/or DCZ treatment (**data not shown**). We attribute this effect to P-TEFb activation driven by JQ1-dependent release of Brd4 from chromatin, a mechanism proposed to account for JQ1-dependent reversal of HIV latency and JQ1-dependent immediate early gene activation in striatal neurons [51,52]. That dBET6 treatment, which depletes Brd4 protein, did not lead to any change in c-myc transcripts, is consistent with this interpretation (**Figure 2B**). Although we did not investigate the mechanism for this effect further, our results are consistent with the notion that JQ1 inhibited BET association with chromatin.

**Figure 2.**
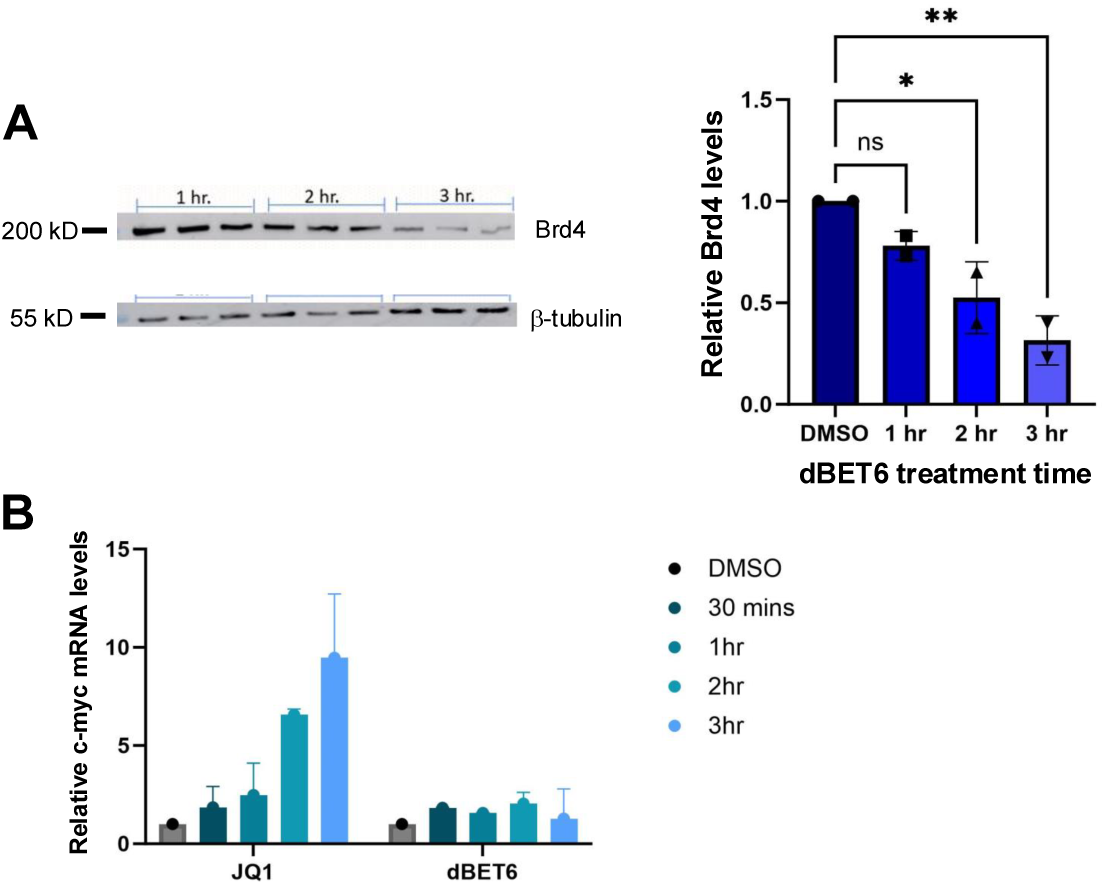
Functional validation of BET inhibitors. **(A)** Left: Representative anti-Brd4 immunoblots of HEK 293 cells treated with 100 nM dBET6 for the indicated times. Right: Plot of ImageJ quantification of anti-Brd4 immunoblots (normalized to loading control, n=3; means +/- SEM). Values for 3-hour vehicle (DMSO) treatment were set to 1. Statistical significance was assessed by one-way ANOVA followed by Dunnet’s post-hoc test; * p<0.1, ** p<0.01. **(B)** RT-qPCR to quantify relative c-myc transcript levels after treatment of HEK 293 cells with either 1 μM JQ1 or 100 nM dBET6 for the indicated times (n=3 independent experiments). Values for 3-hour vehicle (DMSO) treatment were set to 1.

### 3.3 Analysis of G protein activation by RNA sequencing

We then performed RNA-seq in DREADD-expressing cells in the presence or absence of DCZ, and/or BET inhibitor. We chose 1-hour DCZ treatments to capture cells when G protein signaling was still ongoing (see **Figure 1**) and when differentially expressed genes are likely to be detected. These treatment conditions were carried out either alone or in combination with 3-hour inhibitor treatments (such that DCZ was added in the last hour of inhibitor treatment). A control sample that was not treated with DCZ or inhibitor was also included for each DREADD. We analyzed two biological replicates of each condition using RNA-seq. We observed high concordance between replicates as indicated by Pearson correlations and principal component analysis (**Figure S2**).

DCZ treatment significantly increased expression (by 1.5-fold or more; p adj ≤0.1) of 55 genes in cells expressing the Gαq DREADD (relative to the DMSO control) and 4 genes in cells expressing the Gαs DREADD. Interestingly, no transcripts were significantly decreased in either condition (**Figure 3A**). The 55 genes induced by Gαq signaling included a set of 22 genes classified as primary response genes upon ERK activation (**Figure 3B**)[53]. This was expected given that Gαq signaling is known to increase ERK1/2 activation in HEK 293 cells [54]. In addition, KEGG pathway mapping showed significant enrichment of functions related to IL-17, TNF, and NFκB signaling pathways among Gαq-induced genes (**Figure 3C)**.

**Figure 3.**
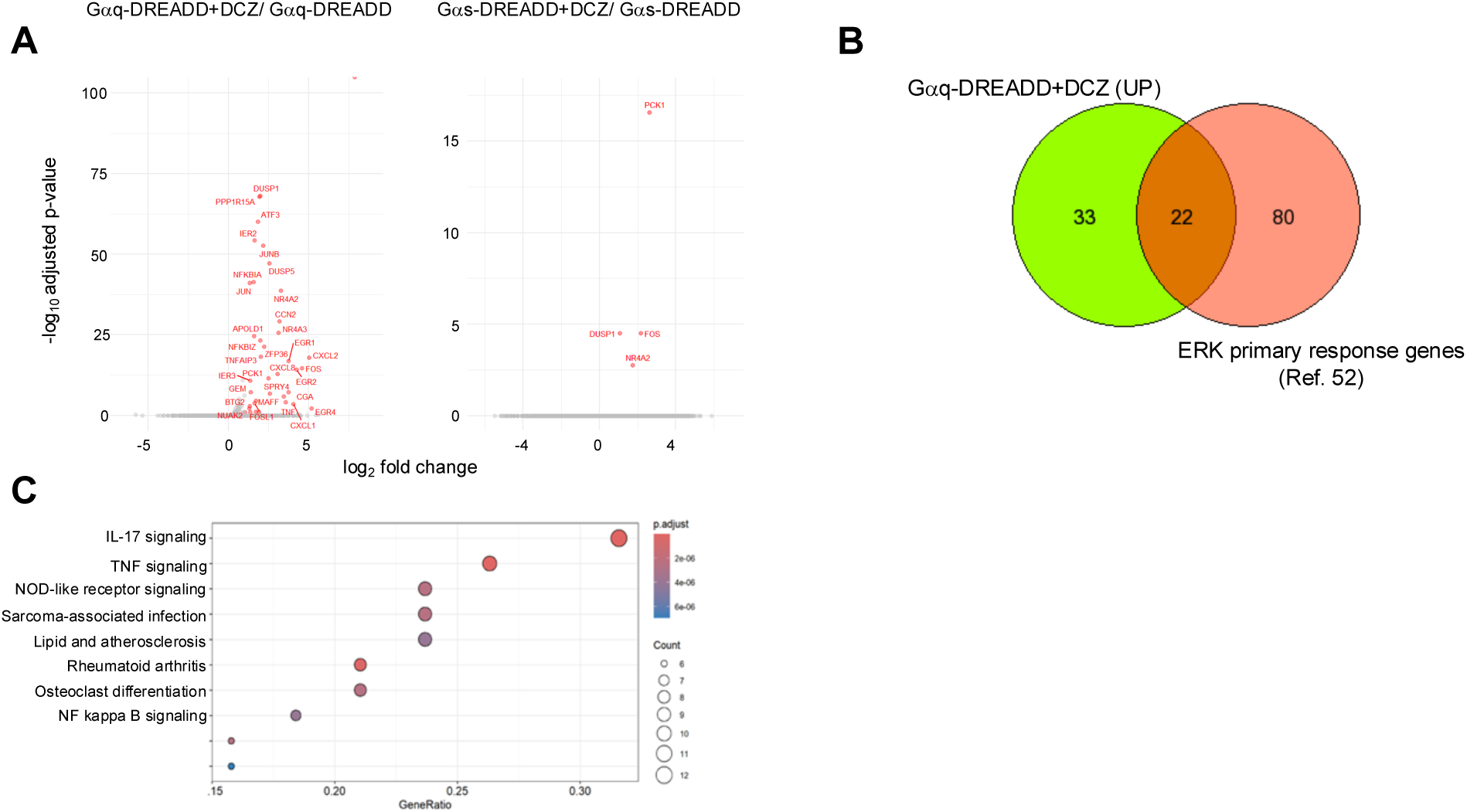
Gene expression changes triggered by DREADDs in HEK 293 cells. **(A)** Left: Volcano plots of differentially expressed genes identified comparing cells expressing the Gαq-DREADD (right) or Gαs-DREADD (left) +/- DCZ treatment (fold change ≥1.5, p≤0.1). **(B)** Venn diagram showing overlap of genes induced by Gαq-DREADD/DCZ and ERK primary response genes identified in [52]. **(C)** KEGG pathway analysis of 55 Gαq-DREADD/DCZ upregulated genes. “GeneRatio” indicates ratio of number of genes enriched in the indicated category to total number of differentially expressed genes.

The four genes (PCK1, DUSP1, FOS, NR4A2) that were significantly upregulated by DCZ in cells expressing the Gαs DREADD were also induced by the Gαq DREADD, and two of them (DUSP1, FOS) are primary response genes (**Figure 3A**)[53]. This suggests a shared function for Gαq and Gαs signaling in activating ERK1/2 in HEK 293 cells.

### 3.4 Effects of BET inhibition on Gαq and Gαs-mediated gene expression

We next tested the impact of BET inhibitors on DREADD-induced gene expression changes. Treatment with dBET6 alone (i.e. no DCZ) decreased levels of 2140 transcripts and increased levels of 552 transcripts in cells expressing the Gαq-DREADD (**Figure 4A)**. Very similar changes were observed in cells expressing the Gαs-DREADD, although the genes upregulated in dBET6-treated cells expressing either DREADD were only ∼50% overlapping (**Figure S3A**). That the DREADDs alone showed slight differences in the presence of dBET6 may indicate that they have subtle effects on signaling in the absence of ligand, as has previously been shown for the Gαs-DREADD in some cell types [41]. Importantly, neither DREADD alone had a significant impact on gene expression in HEK 293 cells, as no differentially expressed genes were identified when comparing the RNA-seq datasets for the two DREADDs in the absence of DCZ (**data not shown**).

**Figure 4.**
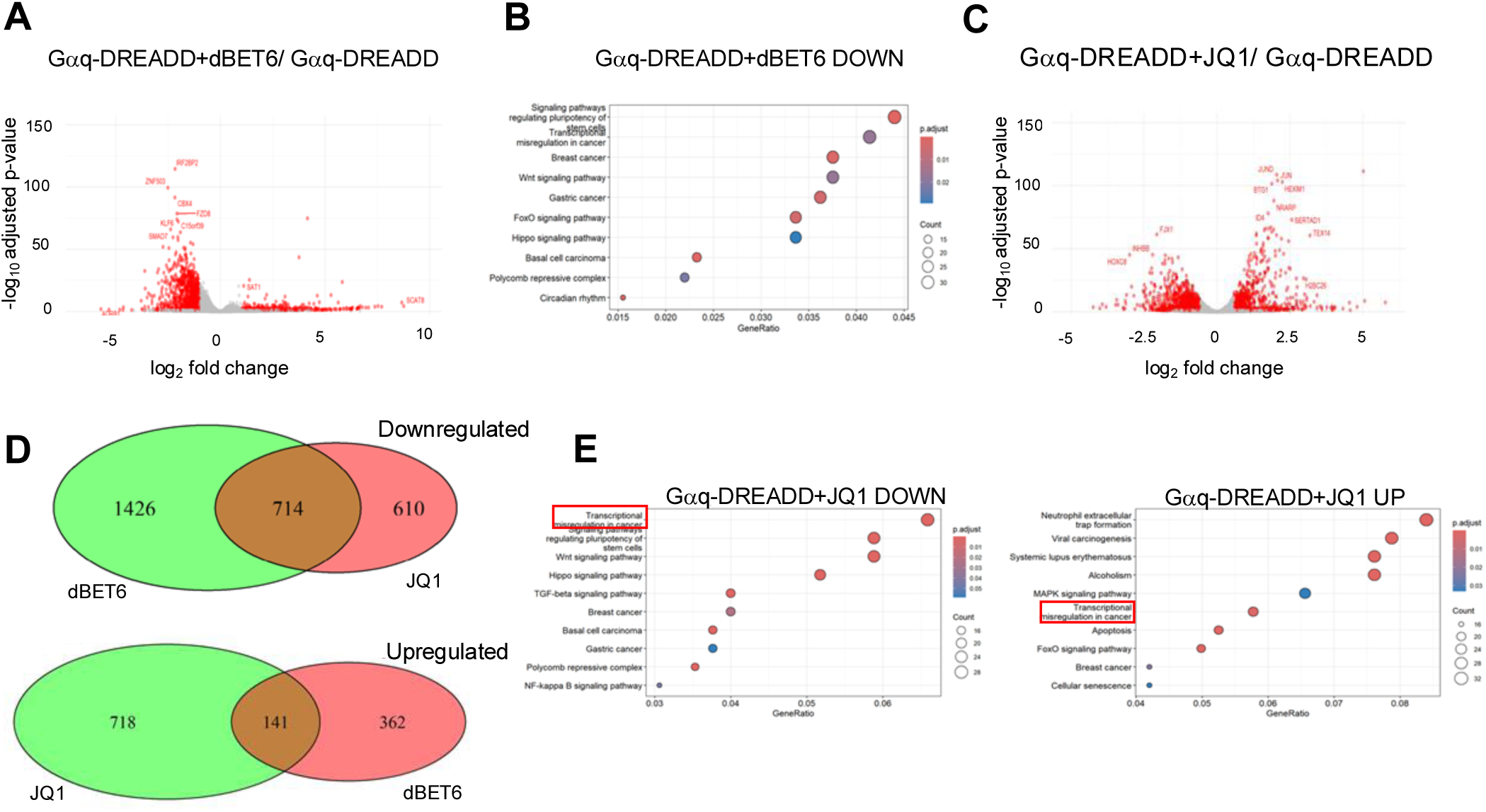
Gene expression changes triggered by BET inhibitors in HEK 293 cells. **(A)** Volcano plot of differentially expressed genes identified comparing cells expressing the Gαq-DREADD +/- dBET6 (fold change ≥1.5, p≤0.1). **(B)** KEGG pathway analysis of significantly downregulated genes from (A). **(C)** Volcano plot of differentially expressed genes identified comparing cells expressing the Gαq-DREADD +/- JQ1 (fold change ≥1.5, p≤0.1). **(D)** Venn diagrams comparing the significantly upregulated (left) and downregulated (right) genes identified upon dBET6 or JQ1 treatment. **(E)** KEGG pathway analysis of significantly upregulated (left) and downregulated (right) genes from (C).

The largely inhibitory effect of dBET6 on transcript levels is consistent with the established function of Brd4 and other BET proteins in transcriptional activation [1]. KEGG pathway analysis of the downregulated genes showed significant enrichment of gene groups linked to signaling and transcriptional regulation in cancer; no significant associations were found among the upregulated genes (**Figure 4B**). JQ1 treatment of Gαq-DREADD expressing cells also altered transcript levels for hundreds of genes, with a more even distribution between upregulation and downregulation (1324 and 859, respectively; **Figure 4C**). The changes were very similar in cells expressing the Gαs-DREADD, with downregulated genes again showing a more complete overlap than upregulated genes (**Figure S3B**). The transcripts decreased by JQ1 were a significantly overlapping subset of those decreased by dBET6 (**Figure 4D**; shown are data for cells expressing the Gαq-DREADD) and KEGG pathway analysis showed significant enrichment of same functional classes of genes downregulated by dBET6 **(Figure 4E)**. There was less pronounced overlap between the transcripts upregulated by JQ1 and dBET6 (**Figure 4D**). Interestingly, KEGG pathway analysis of the JQ1-upregulated genes revealed significant enrichment of genes related to “transcriptional misregulation in cancer,” consistent with our observation that c-myc transcript levels were enhanced (a finding that was confirmed in the RNA-seq data; **data not shown**). HEXIM1 was also significantly induced by JQ1 treatment (**Figure 4C**). HEXIM1 encodes a component of the inactive P-TEFb complex induced upon exposure to stresses that mobilize P-TEFb from its inactive to its active state, thus serving as a feedback regulator of these responses. This is consistent with JQ1-dependent Brd4 release from chromatin mobilizing a fraction of the inactive P-TEFb pool to enhance transcription of some genes [51]. The group of transcripts whose levels were increased by JQ1, but not by dBET6, are likely to be regulated by a similar mechanism.

When compared to vehicle-treated DREADD-expressing cells, cells treated with both BET inhibitor and DCZ showed upregulation and downregulation of sets of genes that overlapped significantly with the same sets identified in cells treated with BET inhibitor alone (**Figure S4**). This is consistent with the large gene regulatory effect of BET inhibition compared to that of DREADD activation in this cell line. However, we observed that the combination of DCZ and BET inhibitors consistently led to larger numbers of affected transcripts (up or downregulated) than BET inhibitors alone (**Figure S4**). This was most evident for transcripts showing increased expression in Gαs DREADD-expressing cells: 638 genes were identified upon treatment with DCZ and dBET6, whereas 419 were identified with dBET6 alone (**Figure S4C**). Thus, although DREADD activation by itself affected expression of small numbers of genes, it modulated the larger effects of BET inhibition.

We found that DCZ treatment of Gαq-DREADD expressing cells in the presence of either dBET6 or JQ1 induced expression of most of the genes induced in the absence of BET inhibitors. In the presence of dBET6, DCZ induced expression of 49 genes, 38 of which were also induced by DCZ in the absence of BET inhibitor (**Figure 5A** and **5B**). DCZ treatment did not downregulate any transcripts in this context, as we observed in the absence of BET inhibitors. 39 genes were induced by DCZ in the presence of JQ1, 36 of which responded to DCZ alone (**Figure 5C** and **5D**). Thus, 65-70% of the genes activated by Gαq signaling in HEK 293 cells are insensitive to BET inhibition. KEGG pathway analysis of the groups of genes resistant to JQ1 or dBET6 showed that they were significantly enriched for functional categories related to cytokine signaling, similar to what we observed in the absence of BET inhibitor (**Figure S5A, S5B,** and **Figure 3A**). This indicated that the character of the Gαq gene expression response was not altered by BET inhibitors.

**Figure 5.**
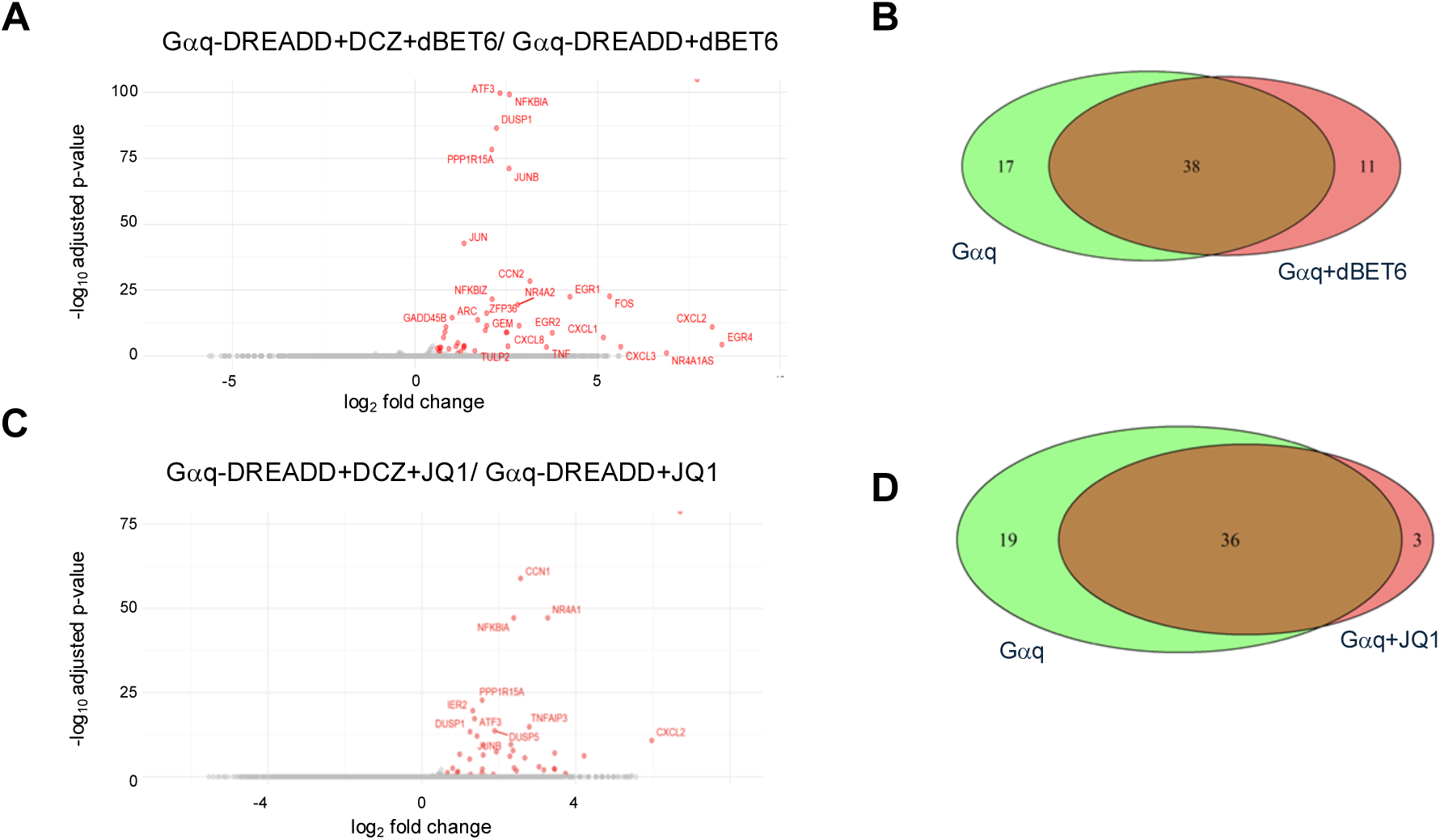
Most gene regulatory effects of Gαq-DREADD activation are maintained in the presence of BET inhibitors. **(A)** Volcano plot of differentially expressed genes identified comparing cells expressing the Gαq-DREADD with combined DCZ/dBET6 treatment to those expressing Gαq-DREADD and treated with dBET6 alone (fold change ≥1.5, p≤0.1). **(B)** Venn diagrams comparing the significantly upregulated genes identified upon DCZ treatment of cells expressing Gαq-DREADD in the presence or absence of dBET6. **(C)** Volcano plot of differentially expressed genes identified comparing cells expressing the Gαq-DREADD with combined DCZ/JQ1 treatment to those expressing Gαq-DREADD and treated with JQ1 alone (fold change ≥1.5, p≤0.1). **(D)** Venn diagrams comparing the significantly upregulated genes identified upon DCZ treatment of cells expressing Gαq-DREADD in the presence or absence of JQ1.

Three of the four genes induced by DCZ treatment of cells expressing the Gαs-DREADD (PCK1, NR4A2, FOS) were not induced in the presence of dBET6 (**Figure S6A; data not shown**). Interestingly, these three genes were among those that were resistant to dBET6 in Gαq-DREADD-expressing cells (**data not shown**). All four genes were induced by DCZ in the presence of JQ1, as well as eight additional genes (**Figure S6B**). The divergent effects of dBET6 and JQ1 in this context may again reflect altered function of BET proteins released from chromatin by JQ1 (see **Figure 2B**). Although based on a small number of genes, these results suggest that Gαq and Gαs signaling led to induction of overlapping sets of genes through distinct mechanisms.

## 4. Discussion

In this study, we expressed DREADDs in HEK 293 cells to specifically assess Gαq- or Gαs-dependent changes in gene expression and assess how BET inhibitors influenced such changes. Although the scope of the gene expression changes associated with DREADD activation in our system was limited, our data point to transcriptional induction by Gαq signaling as largely independent of BET co-regulators. This is consistent with the notion that BET proteins are context-specific regulators that respond to cellular signaling inputs, rather than acting as general transcription factors.

DREADDs have been used to study cellular signaling events extensively in a variety of cell types [48]. To our knowledge, this is the first example of the use of DREADDs to specifically interrogate G protein-mediated changes in gene expression. Stimulation of the Gαq-DREADD in HEK 293 cells activated a subset of immediate early genes. This was consistent with canonical Gαq signaling involving phospholipase Cβ, Ca^2+^ mobilization, protein kinase C (PKC), and subsequent ERK1/2 activation [55,56]. We also observed increased expression of a subset of genes that are normally targets of inflammatory signaling pathways such as IL-17 and TNFα. This subset included a group of linked CXCL chemokine genes co-regulated by a proposed super enhancer [57]. These effects could reflect Gαq-dependent activation of NF-κB through phosphoinositol-3-kinase (PI3K) and Akt-mediated phosphorylation of IκB kinase [58]. Deciphering the nature of pathways preferentially activated by stimulation of the Gαq-DREADD in HEK 293 cells will require further analysis.

Stimulation of the Gαs-DREADD induced expression of only four genes, all of which were among those induced by the Gαq-DREADD. Gαs signaling is also associated with ERK1/2 activation in HEK 293 cells, and so an effect on immediate early genes that overlaps with that of Gαq signaling might be expected [54,55,59]. It is unclear why so few responsive genes were identified upon stimulation of the Gαs-DREADD. We are currently investigating whether this could reflect a rapid burst of gene expression change induced by Gαs signaling that we did not capture in our RNA-seq analysis.

Consistent with previous studies, dBET6 treatment led to more robust effects on steady state transcripts than did JQ1, and its effects were more biased toward reduced transcript levels. This is most likely due to bromodomain-independent functions of BETs, which have been characterized for Brd4 and which are mitigated by dBET6 but not by JQ1 [20,60]. We also observed a group of genes whose transcription was apparently stimulated by the release of BETs from chromatin, including c-myc and HEXIM1. This occurs because Brd4 that is liberated from chromatin by JQ1 is able to engage P-TEFb and stimulate transcription [51]. The effect of JQ1 on immediate early genes that we found here resembles our previous findings in striatal neurons but differs from that reported in cortical neurons [33,52]. Such differences may result from the differing status of cellular signaling pathways between cell types, consistent with the idea that BETs are signal transducers.

In cells expressing the Gαq DREADD, DCZ activated gene expression largely independently of BETs. This was evident from the fact that 65-70% of the DCZ responsive genes were still induced in the presence of either JQ1 or dBET6. Perhaps more striking was the fact that Gαq signaling activated genes linked to cytokine signaling and NFκB even in the presence of BET inhibitors. Inflammatory genes are highly sensitive to BET inhibition in several physiological contexts; the cluster of chemokine genes activated by Gαq signaling in our dataset were previously found to be JQ1-sensitive in liver endothelial cells [3,7,57]. Our previous findings in primary cardiomyocytes indicated that inflammatory genes were insensitive to JQ1 when cells were stimulated with endothelin-1, a ligand that signals through a Gαq-coupled GPCR [38]. The findings here again suggest that Gαq-coupled GPCR signaling can rewire certain transcriptional responses to reduce their dependence on BET protein function, highlighting the sensitivity of BETs to the cellular signaling milieu. The fact that only four genes were induced by DCZ in the presence of the Gαs-DREADD precluded a robust comparison between the gene expression effects of the two signaling pathways. However, all four genes were induced by Gαq signaling as well and in that case, were BET inhibitor-resistant, whereas DCZ-mediated induction of three of the four genes was blocked by dBET6 in the presence of the Gαs-DREADD. This is again reminiscent of our previous findings in cardiomyocytes: when a Gαs-coupled GPCR was stimulated, inflammatory genes were induced in a JQ1-sensitive manner.

BET inhibitor resistance linked to constitutively active JAK2 or mutations in the cullin E3 ligase SPOP is caused by stabilization of Brd4 protein, leading to increased levels that counteract BET inhibition [34,35,37]. We did not observe significant effects of DREADD signaling on Brd4 protein levels in the presence of dBET6, suggesting that Brd4 stabilization does not account for BET inhibitor resistance in our system. Resistance may result from alleviating endogenous BET inhibitory mechanisms, such as binding by the retinoblastoma protein RB [61]. Alternatively, the Gαq DREADD may trigger a BET-independent pathway for inflammatory gene activation, analogous to Wnt pathway activation in BET inhibitor-resistant leukemias [62]. This latter mechanism would be consistent with our previous findings in cardiomyocytes, which showed no change in Brd4 levels upon Gαq-coupled or Gαs-coupled receptor stimulation but differences in Brd4 recruitment to target loci by chromatin immunoprecipitation (ChIP) [38]. Future work will be needed to distinguish between these mechanisms in our system. This will be important in furthering our understanding of the response of BETs to cell signaling pathways and how this can be harnessed in the use of BET inhibitors as therapeutics.

## Acknowledgements

The authors thank Professor Bryan Roth, Department of Pharmacology, University of North Carolina for advice about the use of DREADDs, and members of the Tanny and Hébert labs for helpful feedback and discussions.

## Author contributions

**AJ**: Investigation, Methodology, Data Curation, Visualization; **VP**: Investigation, Methodology; **DD**: Investigation, Methodology, Supervision; **DP**: Investigation, Methodology, Supervision; **TEH**: Conceptualization, Funding Acquisition, Supervision, Writing-review and editing; **JCT**: Conceptualization, Funding Acquisition, Supervision, Writing-original draft, Writing-review and editing.

## Funding sources

This work was supported by the Canadian Institutes for Health Research (PJT-173356 to JCT; PJT-159687 to TEH).

---

**Figure S1.**
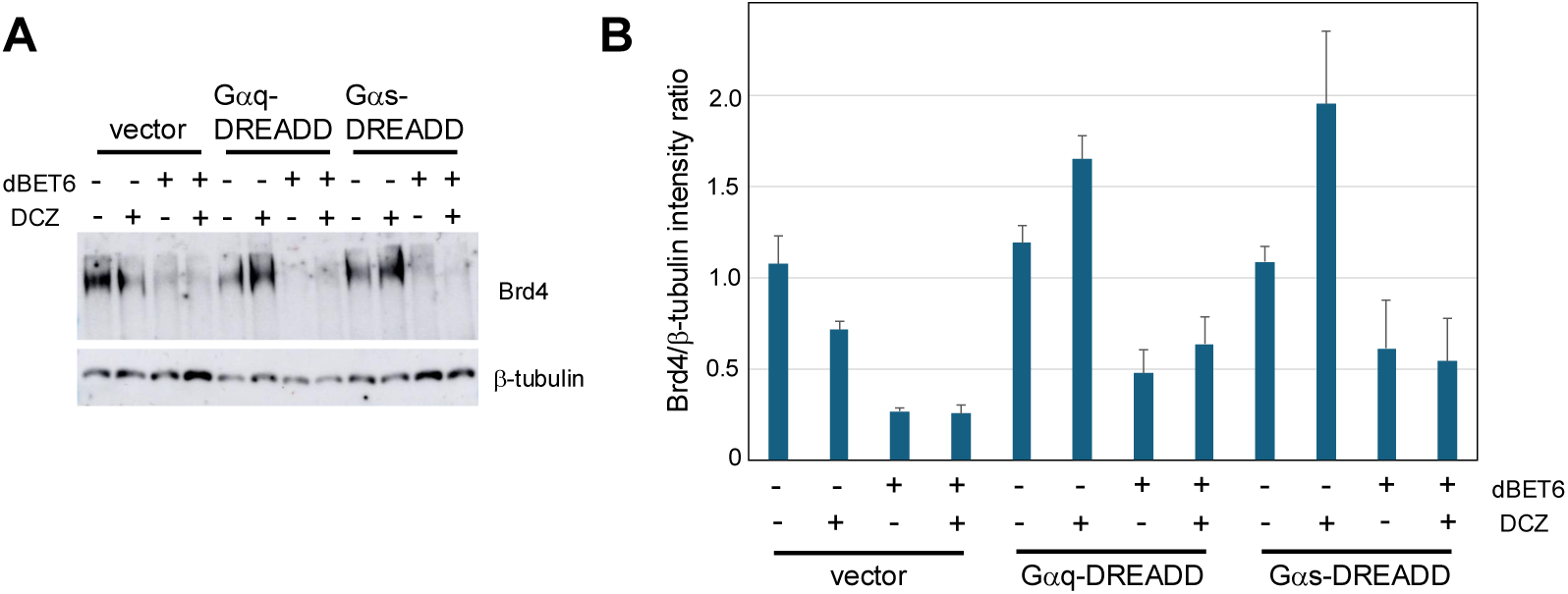
Brd4 depletion by dBET6 in presence and absence of DREADD signaling. **(A)** Representative immunoblot of extracts from HEK 293 cells transfected with the indicated plasmids and treated with dBET6 (100 nM for 3 hours), DCZ (1 μM for 1 hour), or both (with DCZ treatment occurring in the last hour of dBET6 treatment). Antibodies are indicated on the right. **(B)** Quantification of Brd4 signals normalized to β-tubulin under the indicated conditions (n=3; error bars indicate SEM).

**Figure S2.**
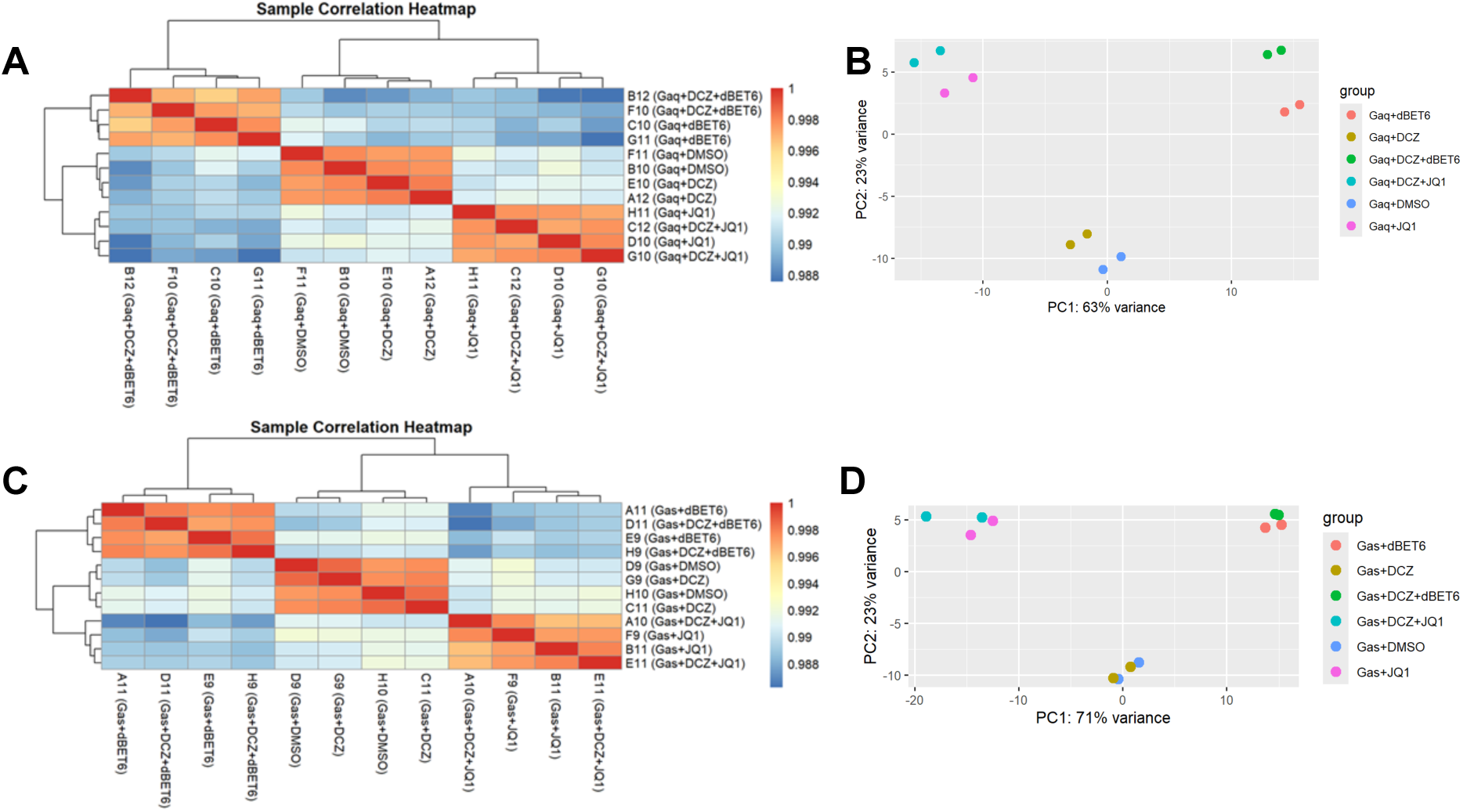
Pearson correlations and principal component analysis for RNA-seq datasets. **(A)** Pearson correlation heatmap for RNA-seq datasets from Gαq-DREADD expressing cells. P-values indicated by the colour bar. **(B)** Principal component analysis of RNA-seq datasets from Gαq-DREADD expressing cells. **(C)** Pearson correlation heatmap for RNA-seq datasets from Gαs-DREADD expressing cells. P-values indicated by the colour bar. **(D)** Principal component analysis of RNA-seq datasets from Gαs-DREADD expressing cells.

**Figure S3.**
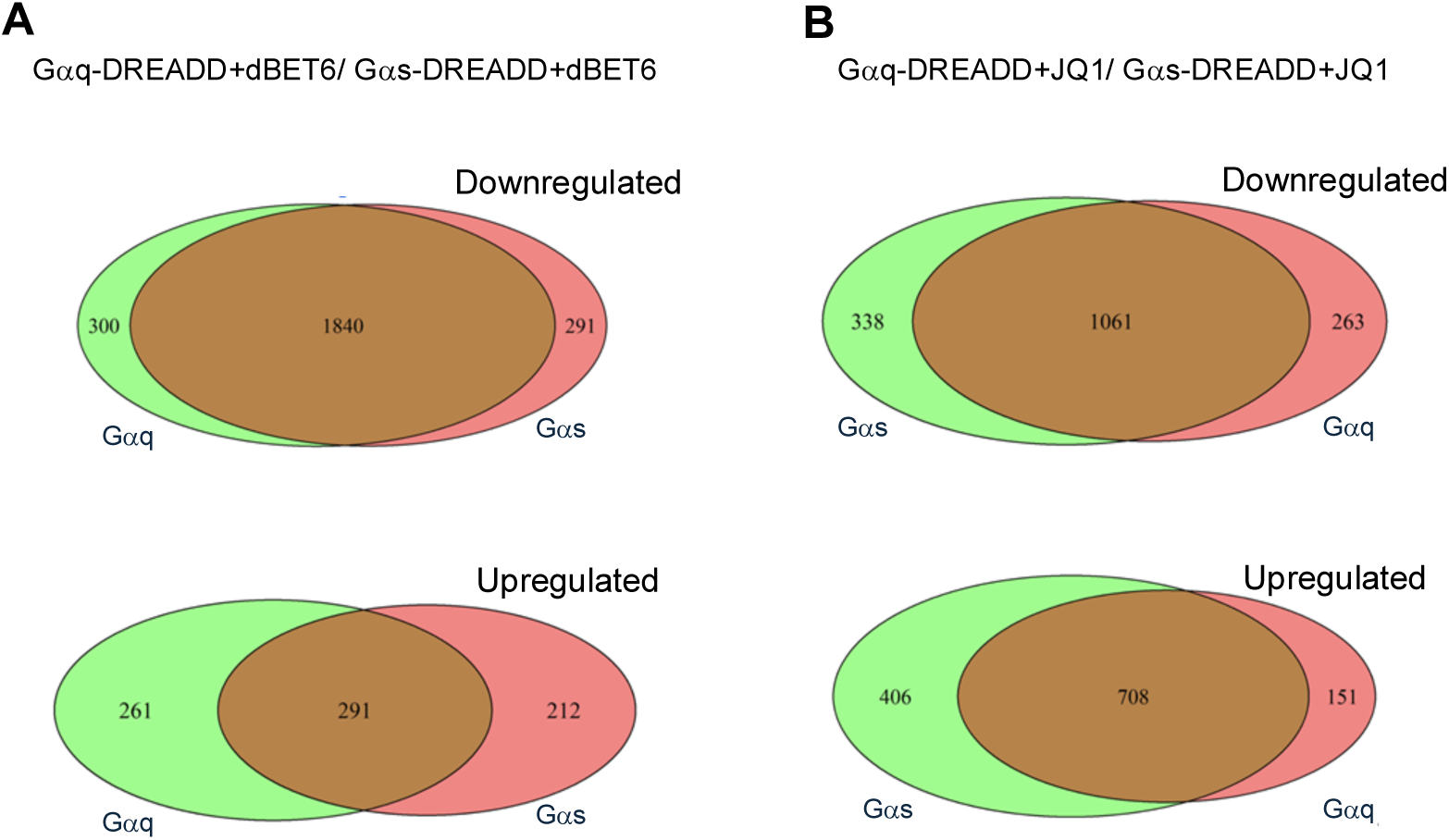
Comparison of the effect of BET inhibitors in cells expressing Gαq-DREADD and Gαs-DREADD in the absence of DCZ. **(A)** Venn diagrams comparing the significantly downregulated (top) or upregulated (bottom) genes identified upon dBET6 treatment of cells expressing Gαq-DREADD or Gαs-DREADD. **(B)** Venn diagrams comparing the significantly downregulated (top) or upregulated (bottom) genes identified upon JQ1 treatment of cells expressing Gαq-DREADD or Gαs-DREADD.

**Figure S4.**
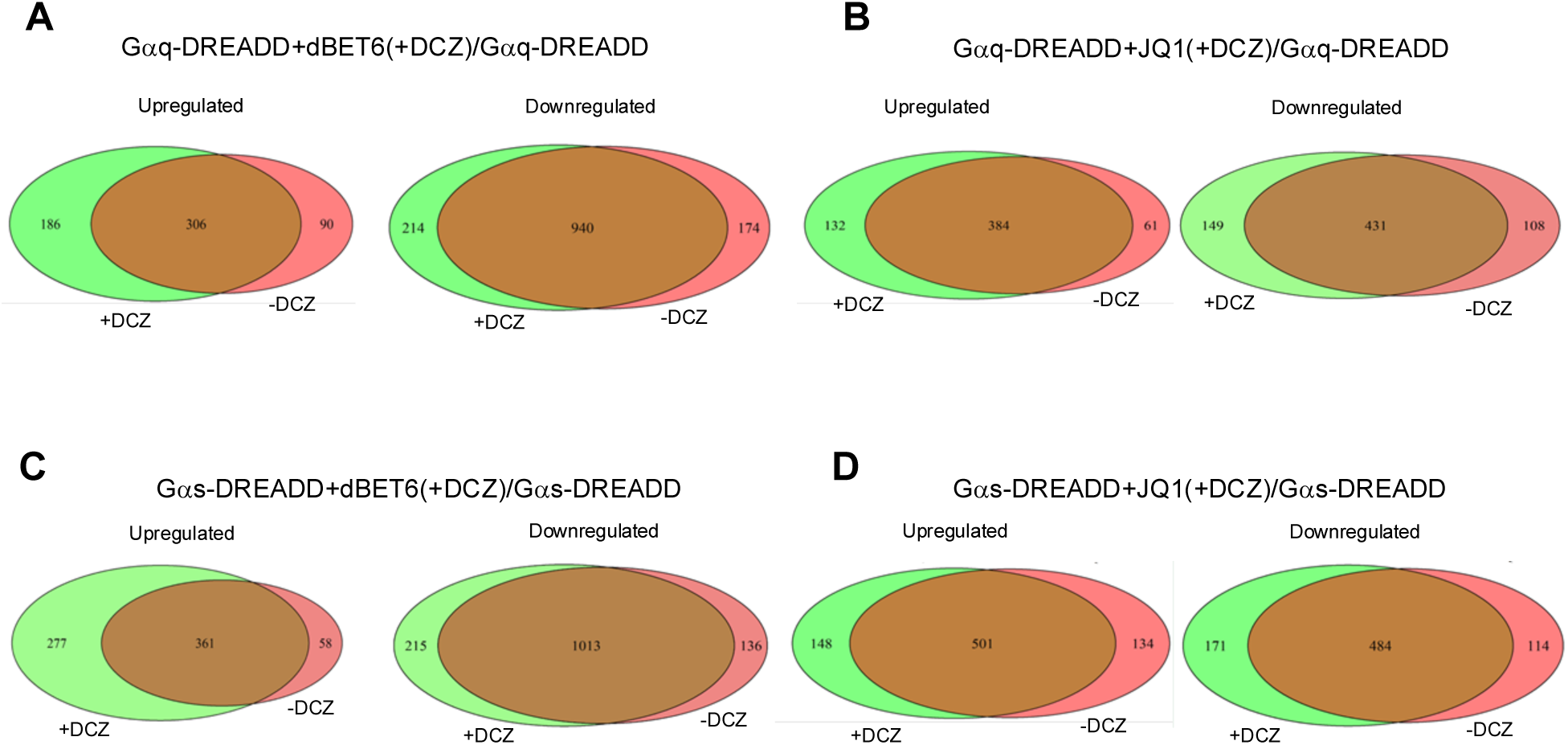
Combined effects of DCZ and BET inhibitors. **(A)** Venn diagrams comparing the significantly upregulated (left) or downregulated (right) genes identified upon combined DCZ/dBET6 treatment of cells expressing Gαq-DREADD. **(B)** Venn diagrams comparing the significantly upregulated (left) or downregulated (right) genes identified upon combined DCZ/JQ1 treatment of cells expressing Gαq-DREADD. **(C)** Venn diagrams comparing the significantly upregulated (left) or downregulated (right) genes identified upon combined DCZ/dBET6 treatment of cells expressing Gαs-DREADD. **(D)** Venn diagrams comparing the significantly upregulated (left) or downregulated (right) genes identified upon combined DCZ/JQ1 treatment of cells expressing Gαs-DREADD.

**Figure S5.**
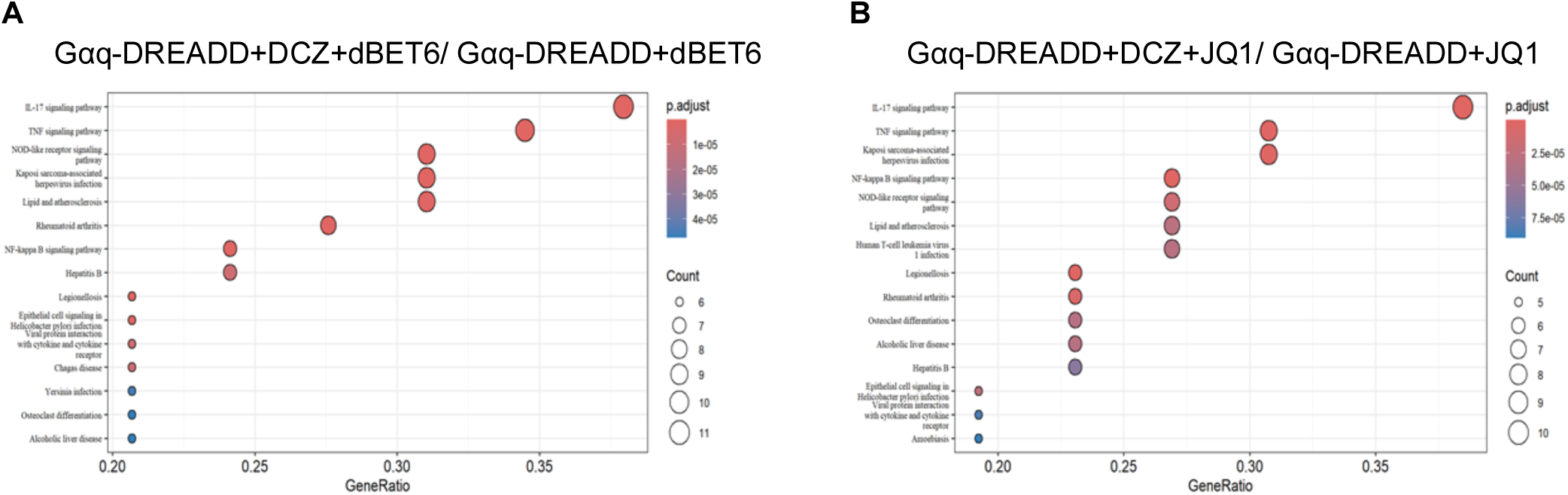
Enrichment of inflammatory functions among genes upregulated by DCZ in Gαq-DREADD expressing cells in the presence of BET inhibitors. **(A)** KEGG pathway analysis of 38 significantly upregulated genes in the presence of dBET6. **(B)** KEGG pathway analysis of 36 significantly upregulated genes in the presence of JQ1.

**Figure S6.**
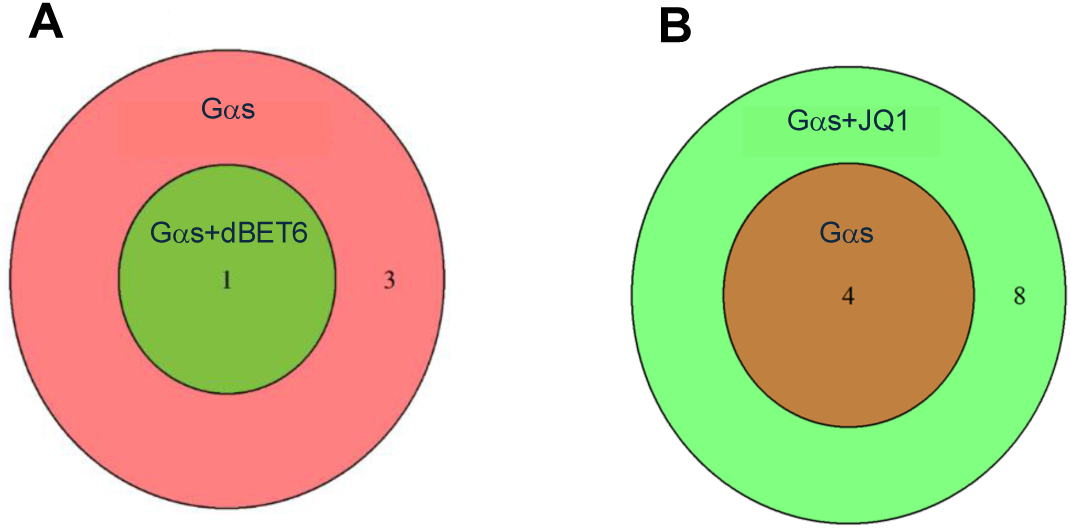
Effects of BET inhibitors on Gαs-induced gene expression. **(A)** Venn diagram comparing significantly upregulated genes identified upon DCZ treatment of cells expressing Gαs-DREADD in the presence or absence of dBET6. **(B)** Venn diagram comparing the significantly upregulated genes identified upon DCZ treatment of cells expressing Gαs-DREADD in the presence or absence of JQ1.

## Notes

### Competing Interest Statement

The authors have declared no competing interest.

